# *Drosophila* larval Odd neurons process innate and learned information to regulate chemotaxis behavior

**DOI:** 10.1101/2025.06.04.657871

**Authors:** Ali Asgar Bohra, Guy Tear, Camilla Larsen, Mikko Juusola

## Abstract

Adaptive decision-making emerges from the integration of innate preferences and learned experiences to guide behavior. Using *Drosophila* larvae as a model, we investigated the neural circuitry underlying olfactory processing, focusing on Odd neurons—a distinct neuronal population that receives innate valence signals from Kenyon cells (KCs) rather than lateral horn inputs. Through larval connectomics, trans-tango labeling, and detailed anatomical analyses, we found that Odd neurons integrate innate valence through dendro-dendritic connections with KCs and learned valence via inhibitory inputs from mushroom body output neurons (MBON-g1 and MBON-g2). Optogenetic silencing of Odd neurons disrupted larval chemotaxis and associative memory by selectively impairing memory retrieval, while sparing memory formation, and by altering the turn rate during navigation. These results suggest that Odd neurons serve as an integrative hub, converging innate olfactory cues with learned reinforcement signals to fine-tune navigational choices. This study illuminates how distinct neural pathways converge in the larval brain to shape behavior, offering novel insights into decision-making circuitry that may extend to more complex nervous systems.

## Introduction

Decision-making processes integrate genetic predispositions and learned experiences to transform complex sensory inputs into unified neural representations that drive adaptive behavior^1,2^. While some responses to sensory stimuli are hard-wired^3^ organisms can adjust their reactions based on prior experiences and environmental changes^4,5^. Remarkably, learning can even reverse the valence, or subjective valence, such as ‘good or bad’, of an innate response^6,7^. However, the precise mechanisms by which innate and learned valences interact remain unclear. This study aims to investigate these mechanisms using the olfactory system of *Drosophila* larvae as a model, capitalizing on its genetic tractability and well-characterized neural circuits.

In adult *Drosophila*, olfactory sensory neurons send their axons to distinct glomeruli in the antennal lobe, where sensory information undergoes initial processing^8^. Projection neurons (PNs) then relay this information to two higher brain centers: the mushroom body and the lateral horn^9^. Several studies suggest that the lateral horn encodes innate valences, whereas the mushroom body predominantly represents learned valences^10,11^.

In *Drosophila* larvae, the components of the mushroom body and their roles in memory formation and retrieval are well characterized^12,13^. Beyond its role in learned valence, several studies suggest that the mushroom body contributes to innate olfactory behavior^14^ (**Figure 1A**). The dendrites of the intrinsic mushroom body neurons—known as Kenyon cells (KCs)— reside in the calyx region, where they receive sparsely encoded olfactory inputs from projection neurons^13^. Their axons form distinct lobe structures that establish synaptic connections with both modulatory dopaminergic neurons (DANs) and lobe-specific mushroom body output neurons (MBONs). DANs deliver the positive or negative reinforcement signals necessary for memory formation^15-17^. When KCs and DANs are simultaneously activated by odor stimuli paired with reward or punishment, synaptic plasticity is induced that modulates MBON responses, ultimately leading to the formation of learned valence. Furthermore, several other intrinsic mushroom body neurons involved in modulatory input, feedback, and convergence also contribute to memory formation and recall^18^.

**Figure 1.**
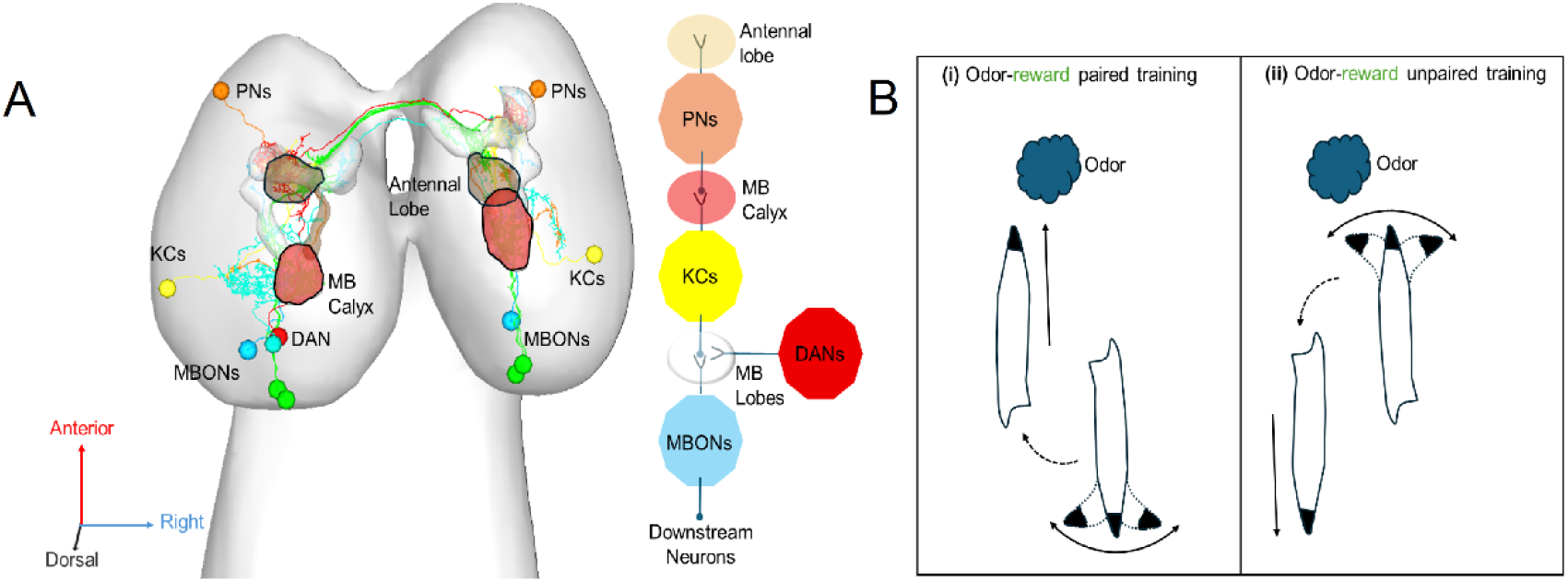
The *Drosophila* larval olfactory learning circuit and chemotaxis following associative learning. (A) Schematic of the *Drosophila* larval olfactory learning circuit. Odor information is initially processed in the Antennal Lobes, where Olfactory Sensory Neurons synapse onto Projection Neurons (PNs). PNs relay this odor information to the Mushroom Body (MB) Calyx, connecting with Kenyon Cells (KCs). Associative learning occurs through modulation by Dopaminergic Neurons (DANs) within the MB Lobes, which convey reward or punishment signals to the KCs. Finally, Mushroom Body Output Neurons (MBONs) integrate the signals from KCs and DANs, driving appropriate behavioural responses via downstream neurons. (B) Larval chemotaxis following odor-based associative learning. (i) Paired odor-reward or unpaired odor-punishment conditioning results in learned attraction, characterized by reduced turning when moving towards the odor source and increased turning when moving away. (ii) Conversely, unpaired odor-reward or paired odor-punishment conditioning results in learned aversion, characterized by increased turning when approaching the odor and reduced turning when moving away.

Recent studies in rodents have highlighted the basolateral amygdala’s role in integrating learned and innate information to produce appropriate behavioral responses^19,20^. Similarly, research on odor perception in *Drosophila* indicates that certain neurons receive innate valence signals directly or indirectly from lateral horn neurons, while learned valence signals are transmitted by neurons downstream of MBONs^11,15^. Recent work has shown that a subtype of convergence neurons (CNs) functionally integrates innate valence from lateral horn neurons with learned valence from MBONs, thereby modulating the larvae’s response to specific odors^21^.

Chemotaxis in *Drosophila* larvae consist of alternating “runs” and turns, guided by run speed, turn rate, and turning direction (**Figure 1B**). While run speed remains largely constant, turn rate and direction are modulated by odor gradients^22^. In innate chemotaxis, the turn rate decreases when odor concentration increases and increases when it decreases^23^. In learned chemotaxis (**Figure 1 Bi**), after odor-reward pairing or odor-punishment pairing, the turn rate decreases when heading toward the odor and increases when moving away. Conversely, after unpaired training (**Figure 1Bii**), the turn rate increases toward the odor and decreases when moving away^24-26^.

In this study, we examine a subset of neurons—termed Odd neurons—that, unlike other local neurons, receive innate valence signals from Kenyon cells rather than lateral horn inputs^27^. Manipulation of Odd neuron activity affects both larval innate chemotaxis behavior and associative memory. We have identified the specific inputs to Odd neurons using larval connectomics and anatomical analyses. Our findings reveal that these neurons receive innate valence via dendro-dendritic connections with Kenyon cells and learned valence via MBONs, specifically MBON-g1 and MBON-g2. This work elucidates how Odd neurons integrate innate and learned valence information to modulate larval chemotaxis behavior.

## Results

### Odd neurons receive information from KCs and MBONs

To elucidate how Odd neurons integrate innate and learned valences, we investigated the neural connections that transmit these signals. Odd neurons comprise a group of 8-10 cholinergic neurons that extend dendrites to the mushroom body’s calyx and project axons to the contralateral centroposterior medial compartment (CPM) (**Figure 2A**). Previous connectomics analyses have suggested that Odd neurons may receive input from Kenyon cells (KCs)^13^. To examine this potential connection, we employed the trans-tango technique— a method for mapping postsynaptic partners in vivo^28^— to label the neurons receiving input from KCs. We used MB247 Gal4 to drive trans-tango expression in KCs^29^ and combined this with Odd LacZ, which selectively marks Odd neurons (**Figure 2B**). Using this approach, we found that 3 out of 8 Odd neurons co-localized with the postsynaptic markers of KCs. This result indicates that these three Odd neurons receive synaptic input from KCs, confirming that KCs are presynaptic to them.

**Figure 2.**
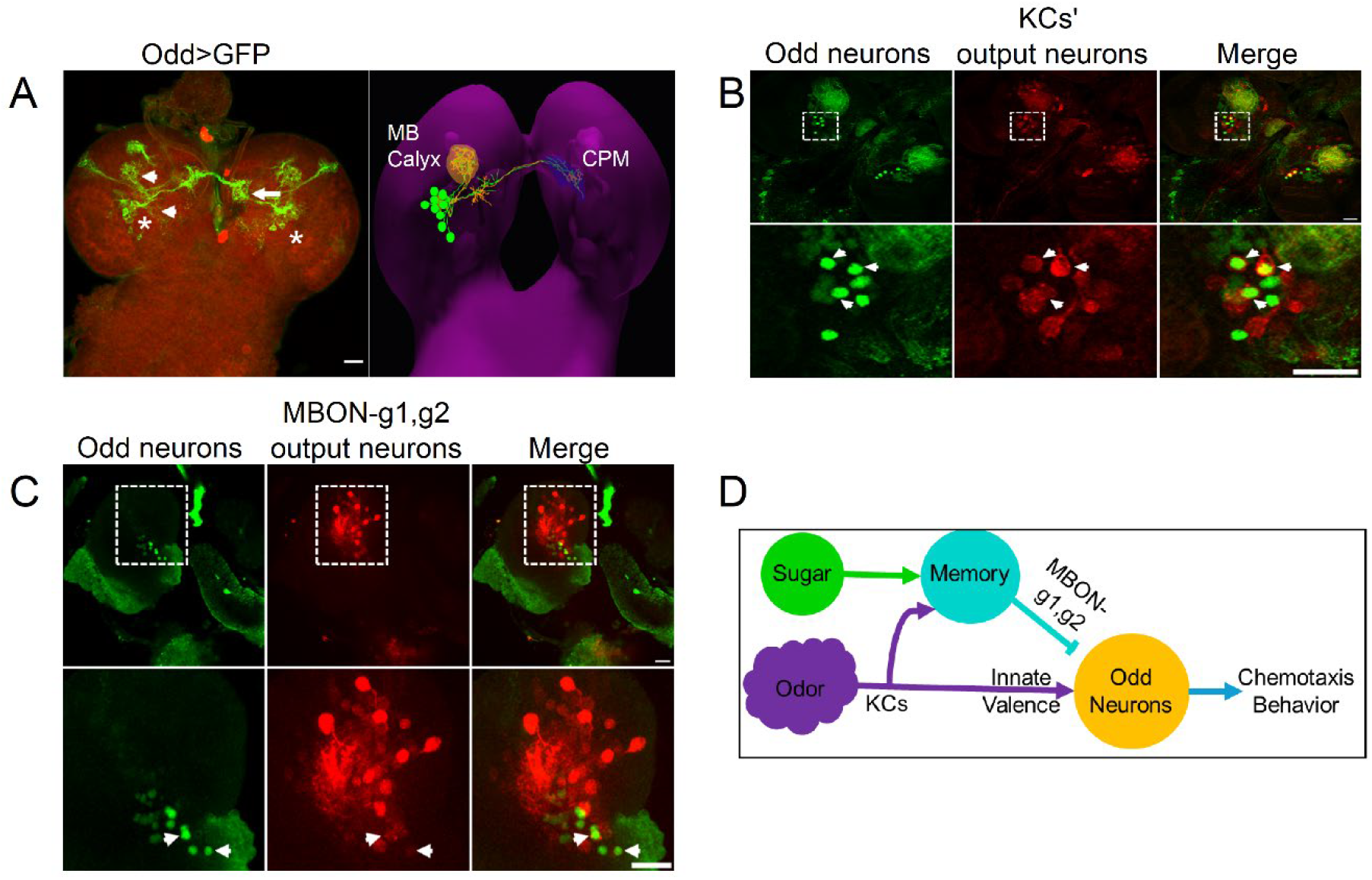
Odd Neurons Integrate Information from Kenyon Cells (KCs) and Mushroom Body Output Neurons (MBONs) (A) Odd-Gal4 labels a group of approximately 8–10 neurons (Odd neurons, indicated by asterisks) characterized by dendritic arborizations in the Mushroom Body (MB) calyx, limited posterior branching, and axonal projections into the contralateral centroposterior medial compartment (CPM). Arrowheads mark dendrites; arrows indicate axonal projections. (B) Trans-Tango labeling identifies three Odd neurons as postsynaptic partners of Kenyon cells (KCs). Odd neurons are shown in green, and KC postsynaptic partners are labeled in red. The arrow highlights the colocalization of Odd neurons with KC output neurons. (C) Trans-Tango labeling reveals two Odd neurons as postsynaptic partners of MBON-g1/g2. Odd neurons are indicated in green, while postsynaptic partners of MBON-g1 and MBON-g2 appear in red. Arrows indicate colocalization. (D) Diagram illustrating the proposed neural circuit where Odd neurons integrate innate and learned valence. KCs transmit innate valence signals to Odd neurons via dendro-dendritic synapses and encode conditioned stimuli essential for associative memory formation. MBON-g1 and MBON-g2 convey learned valence signals to Odd neurons. Thus, Odd neurons serve as convergence points, integrating innate and learned valence inputs to guide chemotactic behavior. Scale bar = 20 µm.

Several studies have suggested that Odd neurons include two specific mushroom body output neurons: MBON-a1 and MBON-a2^30,31^. To confirm this, we conducted a co-localization experiment using RFP to label MBON-a1 and a2 (driven by 52E12 Gal4) and Odd LacZ to mark Odd neurons. Our findings revealed co-localization between MBON-a1/a2 and Odd neurons (**Supplementary Figure 1**), supporting the idea that MBON-a1 and MBON-a2 are indeed part of the Odd neuron population.

Data from larval connectomics indicate that MBON-a1 and MBON-a2 receive inputs from KCs and MBON-g1, MBON-g2, and other neurons^13^. Based on these observations, we propose that KCs, along with other neurons, convey innate valence signals to Odd neurons, while MBON-g1, MBON-g2, and other neurons transmit learned valence to Odd neurons.

Therefore, we further examined the connection between odd neurons and MBON-g1/g2 using trans-tango (**Figure 2C**). By employing R21D06 Gal4 to express trans-tango in MBON-g1 and MBON-g2 and combining it with Odd LacZ, we discovered that two Odd neurons co-localized with postsynaptic markers of MBON-g1 and MBON-g2. This suggests that Odd neurons also receive synaptic input from MBON-g1 and MBON-g2.

### Odd Neurons are Involved in the Retrieval of Associative Memory

*Drosophila* larvae can learn to associate an odor with either a reward or a punishment. To assess their learning ability, we measured the larvae’s odor preference after the odor was paired with a reward (**Figure 3A**). In our experiments, larvae were trained either with an odor paired with a fructose reward or with an odor presented without fructose, with the training repeated twice. Immediately after training, the larvae were divided into two groups: one tested with fructose, the other without, and a performance index was calculated to quantify the extent of their associative olfactory learning.

**Figure 3.**
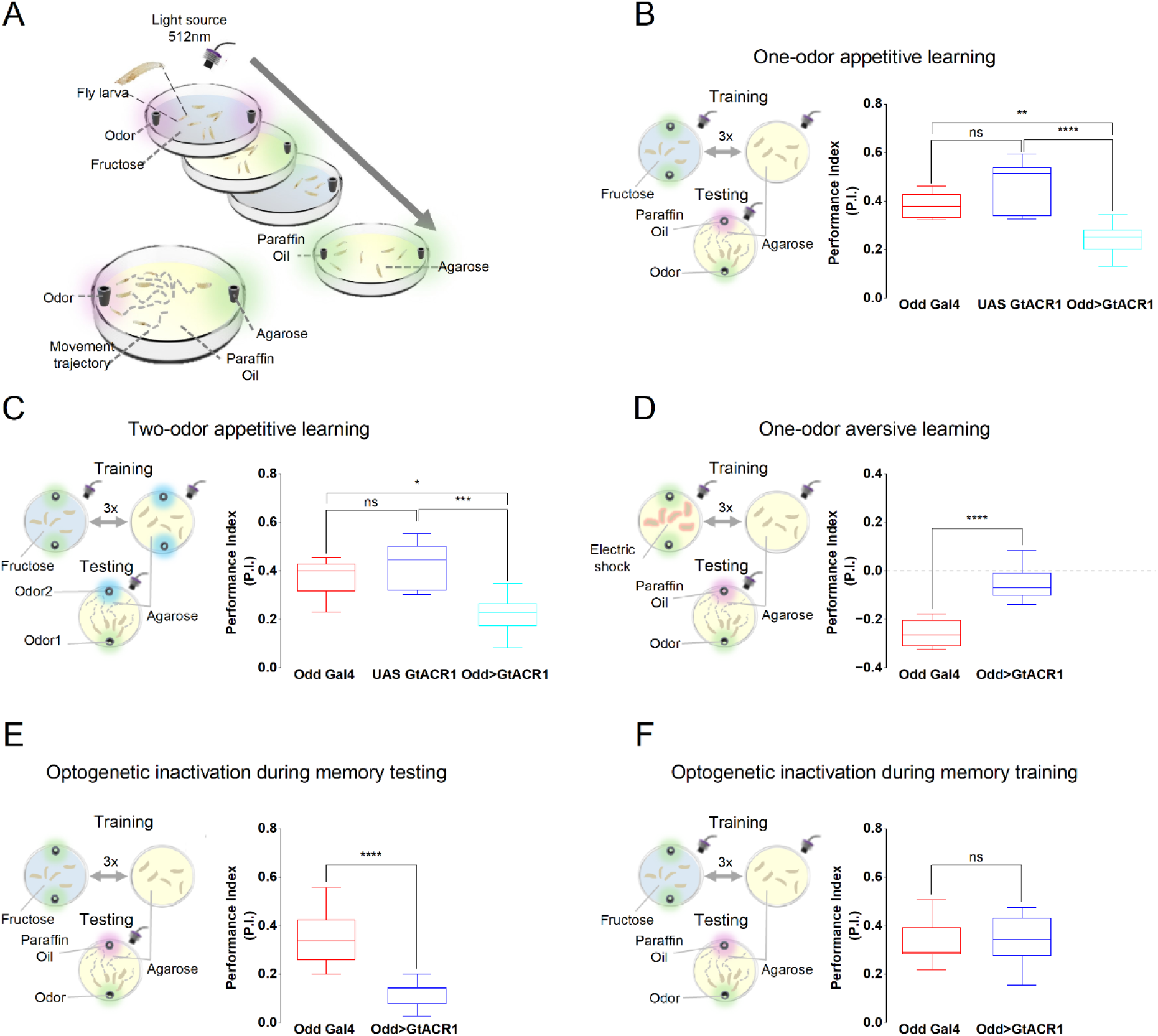
Odd Neurons Are Involved in the Retrieval of Associative Memory. (A) Schematic illustration of the larval olfactory conditioning assay. *Drosophila* larvae were trained through sequential exposure to an odor paired with fructose reward for 5 minutes, followed by exposure to paraffin oil (or vice versa). During testing, larvae were placed on an agarose plate, with the previously trained odor on one side and paraffin oil on the opposite side. Performance Index (P.I.) was determined by counting larvae on each side after 3 minutes. Optogenetic inactivation was conducted using a 512 nm LED when indicated. (**B–F**) Box plots illustrating Performance Index (P.I.) scores of larvae expressing GtACR1 in Odd neurons (Odd>GtACR1) compared to genetic controls (Odd-Gal4 and/or UAS-GtACR1). Schematics alongside each panel illustrate corresponding training and testing procedures. (B) One-odor appetitive learning. Optogenetic inactivation of Odd neurons (Odd>GtACR1) significantly impaired associative learning when odor (AM) was paired with fructose reward, compared to genetic controls (UAS-GtACR1 and Odd-Gal4). (**p < 0.01, ****p < 0.0001, ns = not significant, N ≥ 10). (C) Two-odor appetitive learning. Larvae trained with two odors (odor 1: AM paired with fructose; odor 2: OCT paired with agarose). Optogenetic inactivation of Odd neurons significantly impaired appetitive learning, relative to controls. (*p < 0.05, ***p < 0.001, ns = not significant, N ≥ 10). (D) One-odor aversive learning. Larvae received odor paired with electric shock. Odd neuron inactivation significantly impaired aversive learning compared to control (****p < 0.0001, N ≥ 10). (E)Optogenetic inactivation during memory testing. Larvae expressing Odd>GtACR1 exhibited significantly reduced performance compared to control (****p < 0.0001, N ≥ 10). (F) Optogenetic inactivation during memory training. No significant difference between Odd>GtACR1 and control larvae (ns = not significant, N ≥ 10).

We investigated the role of Odd neurons in learning and memory by inactivating them using the anionic channelrhodopsin GtACR1-activated by a 512 nm green light^32^-during an appetitive associative learning assay (**Figure 3B**). When we paired the odor AM with a fructose reward while inactivating Odd neurons, we observed a significant decrease in the performance index compared to the control group, indicating that Odd neurons contribute to associative memory and learning. Similar results were obtained when two odors (AM and OCT) were used as conditioned stimuli (**Figure 3C**). Because both genetic control lines—Odd-Gal4 and UAS-GtACR1—exhibit similar behavioral phenotypes, we used them interchangeably as controls in subsequent experiments. Interestingly, a decrease in the performance index was also evident during aversive associative memory when the odor AM was paired with an electric shock as punishment (**Figure 3D**). These findings suggest that Odd neurons play a role in appetitive and aversive associative learning.

To further elucidate the impact of Odd neurons on learning and memory, we silenced them at different stages of the training and testing processes. This approach allowed us to pinpoint their specific involvement in the learning and memory cycle.

When Odd neurons were silenced only during testing, we observed a decrease in the performance index (**Figure 3E**). In contrast, silencing them solely during training did not affect the performance index (**Figure 3F**). These results indicate that Odd neurons are crucial for associative learning, particularly for memory retrieval rather than memory formation.

### Odd Neurons Process Both Innate and Learned Valence

Silencing Odd neurons reduces olfactory chemotaxis in naïve larvae^27^. We inactivated Odd neurons optogenetically using the anionic channelrhodopsin GtACR1 and performed a chemotaxis assay (**Figure 4A**). Compared to control larvae, inactivation of Odd neurons resulted in a decreased odor preference, consistent with previous findings (**Figure 4A**). Notably, silencing Odd neurons affects both innate chemotaxis and learned behavior.

**Figure 4.**
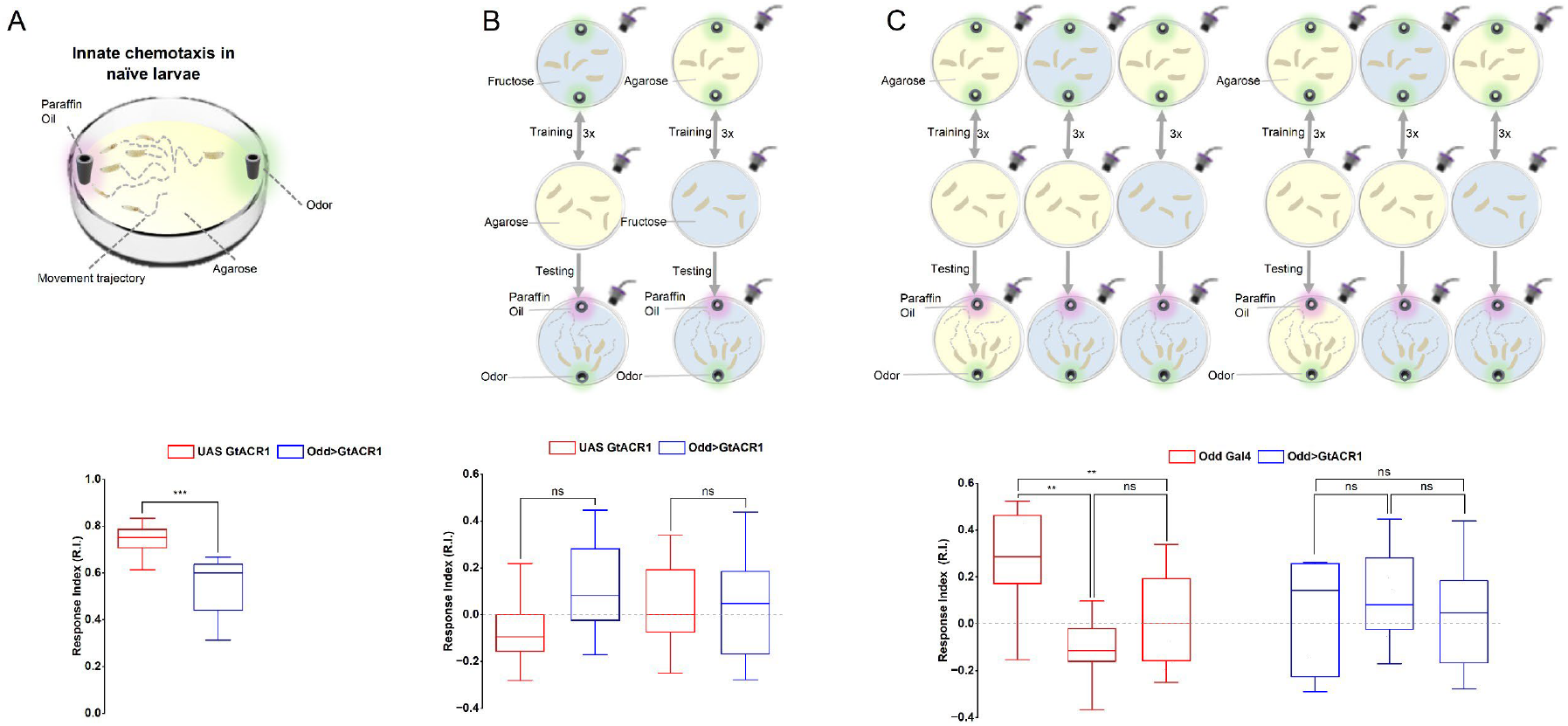
Odd Neurons Process Both Innate and Learned Valence. (A) Optogenetic inactivation of Odd neurons (Odd>GtACR1) significantly reduced innate odor preference in naïve larvae compared to control larvae expressing only UAS-GtACR1 (**p < 0.001). (B) Baseline odor preference in larvae trained with odor-fructose associative memory did not differ significantly between control larvae (UAS-GtACR1) and larvae with optogenetically inactivated Odd neurons (Odd>GtACR1). Similar results were observed in both paired (odor associated with fructose) and unpaired (odor and fructose presented separately) training conditions (ns = not significant). (C) Baseline odor preference after odor-reward associative learning was significantly lower in compared to innate odor preference in control larvae (Odd Gal4) pre-exposed to the odor (**p < 0.01). In contrast, no significant difference was observed in larvae with optogenetically inactivated Odd neurons (Odd>GtACR1) (ns = not significant). *Sample sizes*: N ≥ 10 experiments, each with 30 larvae.

To determine whether the observed behavioral changes during associative learning stem from impaired innate chemotaxis or impaired learned behavior, we conducted associative memory tests in the presence of a fructose reward. The rationale behind this experiment is that the fructose reward should abolish any associative memory while leaving innate olfactory behavior unchanged, thereby reflecting the baseline odor preference^24^.

If the observed behavioral changes are due to impaired learned behavior, the baseline odor preference would remain unchanged between larvae with inactivated Odd neurons and control larvae. Conversely, if the changes resulted from impaired innate olfactory behavior, the two groups’ baseline odor preferences would differ. Our associative memory test with the fructose reward (**Figure 4B**) revealed no significant difference in baseline odor preference between larvae with inactivated Odd neurons and the control group. This finding indicates that the behavioral phenotype observed during associative learning is attributable to impaired learned valence rather than alterations in innate olfactory processing.

We further hypothesized that if Odd neurons mediate learning-associated behavioral changes, then there should be no difference in innate odor preference before and after associative learning in larvae with inactivated Odd neurons. To test this, we performed an appetitive associative learning assay. We compared the baseline odor preference of larvae with silenced Odd neurons to that of larvae exposed to the same odor during the training period. Since odor exposure during training can modify preference relative to naïve larvae, we used the odor preference of pre-exposed larvae as a reference.

Our findings revealed no significant difference in odor preference between learned and pre-exposed larvae with inactivated Odd neurons (**Figure 4C**). In contrast, when Odd neurons were not silenced, a significant difference was observed in baseline odor preference between learned and odor-exposed larvae.

Together, these results suggest that Odd neurons specifically mediate the behavioral changes associated with learning without altering innate olfactory responses.

### Odd Neurons Modulate Turn Rate during Chemotaxis

Larval chemotaxis is characterized by alternating runs and turns^33^. Typically, larvae maintain a straight course with fewer turns when moving toward an odor source and increase their turn rate when moving away to reorient toward the source^23,33^ (**Figure 1Bi**). Following associative training with an odor paired with fructose, larvae display a decreased turn rate when approaching the odor and an increased turn rate when moving away. In contrast, after unpaired training, the pattern reverses: larvae exhibit an increased turn rate when moving toward the odor and a decreased turn rate when moving away^24^. These observations led us to investigate the role of Odd neurons in modulating turn rate during chemotaxis.

When we silenced Odd neurons during an appetitive associative memory test, significant changes in turn rate were observed in both paired and unpaired larvae. Specifically, in the paired condition (**Figure 5A**), larvae with inactivated Odd neurons exhibited an increased turn rate when moving toward the odor source, whereas unpaired larvae showed a decreased turn rate compared to controls. Conversely, paired larvae demonstrated a decreased turn rate when moving away from the odor (**Figure 5B**), while unpaired larvae exhibited an increased turn rate relative to the control group. Notably, the overall total turn rate remained unchanged by the inactivation of Odd neurons.

**Figure 5.**
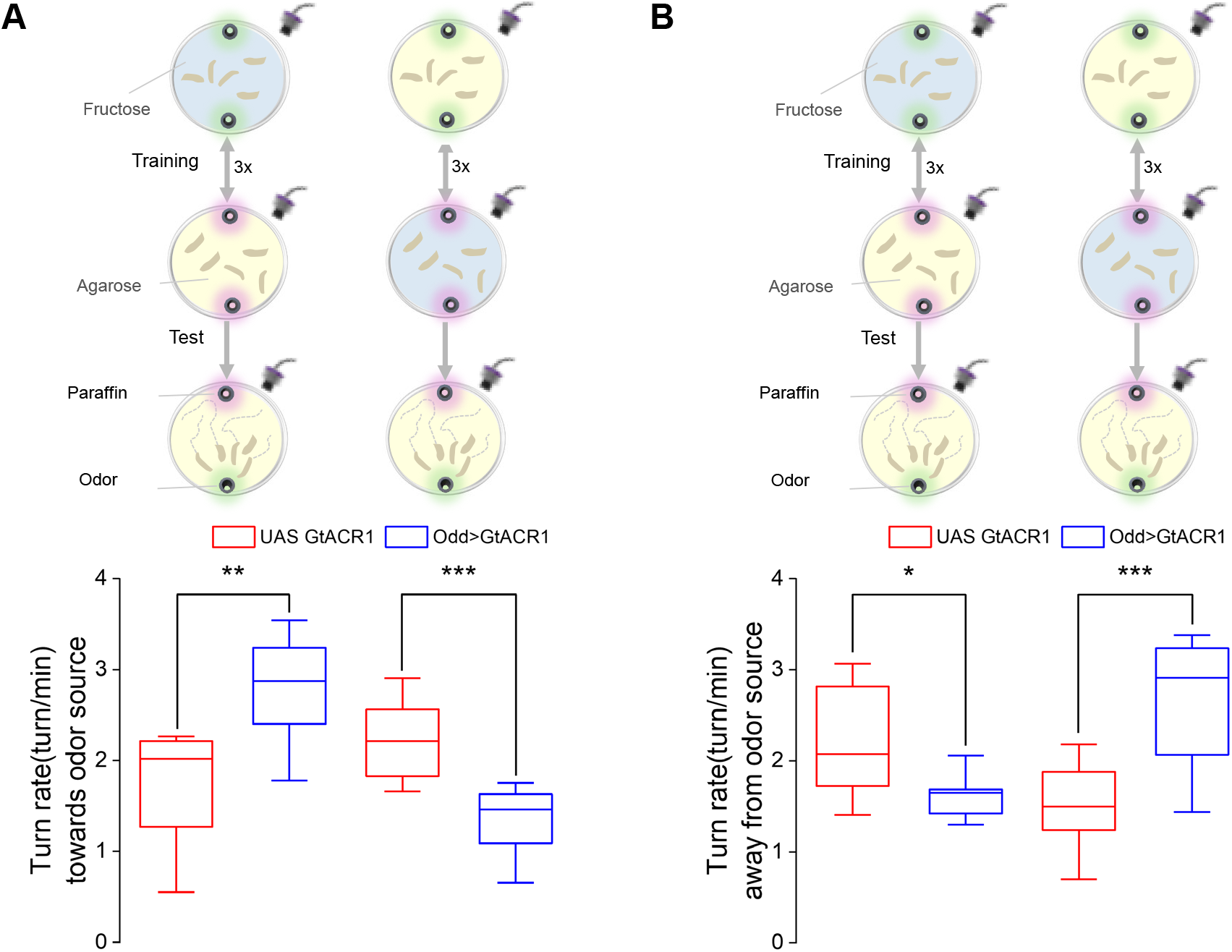
Odd Neurons Modulate Turning behavior during Chemotaxis. (A) Optogenetic inactivation of Odd neurons (Odd>GtACR1) significantly alters larval turning towards the odor source after odor-reward associative learning. Larvae exhibited increased turning towards the odor following paired training (**p < 0.01) and decreased turning following unpaired training (***p < 0.001), relative to control larvae (UAS-GtACR1). (B) Optogenetic inactivation of Odd neurons (Odd>GtACR1) significantly alters larval turning away from the odor source after odor-reward associative learning. Larvae showed decreased turning away from the odor following paired training (*p < 0.05) and increased turning after unpaired training (***p < 0.001), compared to control larvae (UAS-GtACR1). Sample sizes: N ≥ 10 experiments, with 10 larvae per experiment set.

Additionally, when Odd neurons were silenced during appetitive associative memory testing on a fructose plate (**Supplementary Figure 2**), larval turn rates showed no change, regardless of movement direction, compared to odor-exposed larvae.

Together, these results suggest that Odd neurons are essential for fine-tuning turn rate during learned chemotaxis. They adjust the turning behavior as larvae navigate toward or away from an odor source.

## Discussion

In this study, we investigated the role of a specific group of neurons—termed Odd neurons— that integrate innate and learned valence information to regulate larval chemotaxis behavior. Previous work has identified several neurons that integrate innate and learned valence from lateral horn and mushroom body circuits, respectively^13,15,21^. However, our study focuses on Odd neurons, which receive innate valence signals from Kenyon cells (KCs) rather than from lateral horns.

Odd neurons have been implicated in chemotaxis and shown to exhibit GRASP connectivity with projection neurons (PNs) and Kenyon cells^27^. Based on these findings, we hypothesized that Odd neurons receive both innate and learned valence signals. However, our trans-Tango data, together with a recent study, suggest that Odd neurons lack synaptic connection with PNs^31^ while receiving inputs from KCs (**Figure 2B**). Furthermore, previous studies and connectomic analyses indicate that Odd neurons include specific mushroom body output neurons (MBONs)—namely MBON-a1 and MBON-a2—which receive input from KCs as well as from MBON-g1, g2^13,15^. In addition, our trans-tango analysis revealed that Odd neurons receive synaptic inputs from MBON-g1 and MBON-g2 (**Figure 2C**), suggesting that they integrate innate valence from KCs and learned valence from these MBONs. It would be interesting to investigate further the postsynaptic partners of Odd neurons and MBON-a1/a2 and their roles in processing valence. Preliminary trans-tango data indicate that MBON-a1 and a2 neurons form five or more postsynaptic connections, although the identities of these neurons remain unverified.

Our associative memory experiments and innate chemotaxis assay suggest that Odd neurons play a vital role in integrating learned behavior with innate responses (**Figure 3** and **Figure 4A**). The baseline preference experiment using a fructose test plate conclusively shows that the behavioral phenotype observed in larvae with inactivated odd neurons is due to impaired learning rather than altered innate chemotaxis.

*Drosophila* larval chemotaxis relies on the modulation of turning and crawling in response to changing odor concentration^23^. Learned valence further modulates turn rate, steering navigation towards or away from odors. In our study, inactivation of Odd neurons led to an increased turn when larvae moved toward an appetitive odor and a decreased turn rate when moving away (**Figure 5**), suggesting a role for these neurons in memory retrieval for proper navigation. Moreover, when memory expression was abolished by the presence of reward during testing, turn rates did not significantly differ from those of odor-pre-exposed larvae.

Previous studies have shown that inactivating odd neurons increases the turn rate in naïve larvae^27^. However, in our experiments using a fructose test plate, larvae with inactivated Odd neurons did not exhibit a significant change in turn rate compared to control and odor-exposed larvae, suggesting that odd neurons specifically modulate turn rate and chemotaxis associated with learned valence.

Odd neurons appear to serve as the central site for integrating innate and learned valence in the previously proposed model^24^. In this framework, Odd neurons act as the summation point, receiving innate valence input from Kenyon cells (KCs). The GABAergic MBON-g1 and MBON-g2 likely provide inhibitory learned valence, while additional, yet unidentified neurons may contribute excitatory learned valence. Identifying these other presynaptic partners of Odd neurons will be essential for fully understanding how this circuit computes and balances valence to guide behavior.

Overall, our study identifies a specific pathway within the mushroom body in which Odd neurons integrate innate valence signals from KCs with learned valence signals mediated by MBONs, thereby refining chemotaxis behavior. By mapping this circuit, we shed light on how innate and learned information converge to shape decision-making in larval *Drosophila*.

Future studies should determine the postsynaptic targets of MBON-a1 and MBON-a2 to elucidate how valence-integrated signals influence downstream motor circuits. Additionally, further investigation is needed to understand how innate valence circuits from the lateral horn interact with those processed within the mushroom body, particularly in the context of learned experience. Such studies will advance our understanding of how different innate and learned valence pathways converge and coordinate to produce appropriate behaviors across species.

In summary, Odd neurons represent a crucial node in the larval mushroom body, bridging innate and learned valence to regulate navigation. These findings refine our understanding of the functional architecture of the mushroom body and lay the groundwork for further exploration of the neural circuits underlying behavior in *Drosophila*.

## Acknowledgments

We thank Michael Schleyer, Adil Khan and Darren Williams for productive discussions and support. We thank H. MaBouDi, J. Takalo, A. Bridges, and J. Kemppainen for fruitful discussions. This work was supported by BBSRC (BB/P019676/1) to C.L. and G.T. and BBSRC (BB/F012071/1 and BB/X006247/1), EPSRC (EP/P006094/1 and EP/X019705/1), and Leverhulme (RPG-2024-016) grants to M.J.

## Author contributions

A.A.B.: conceptualization, methodology, investigation, formal analysis, visualization, writing (original draft preparation).

C.L.: funding acquisition, conceptualization, supervision, resources, writing (review and editing).

M.J.: funding acquisition, supervision, resources, writing (review and editing) G.T.: supervision, resources, writing (review and editing)

## STAR⋆Methods

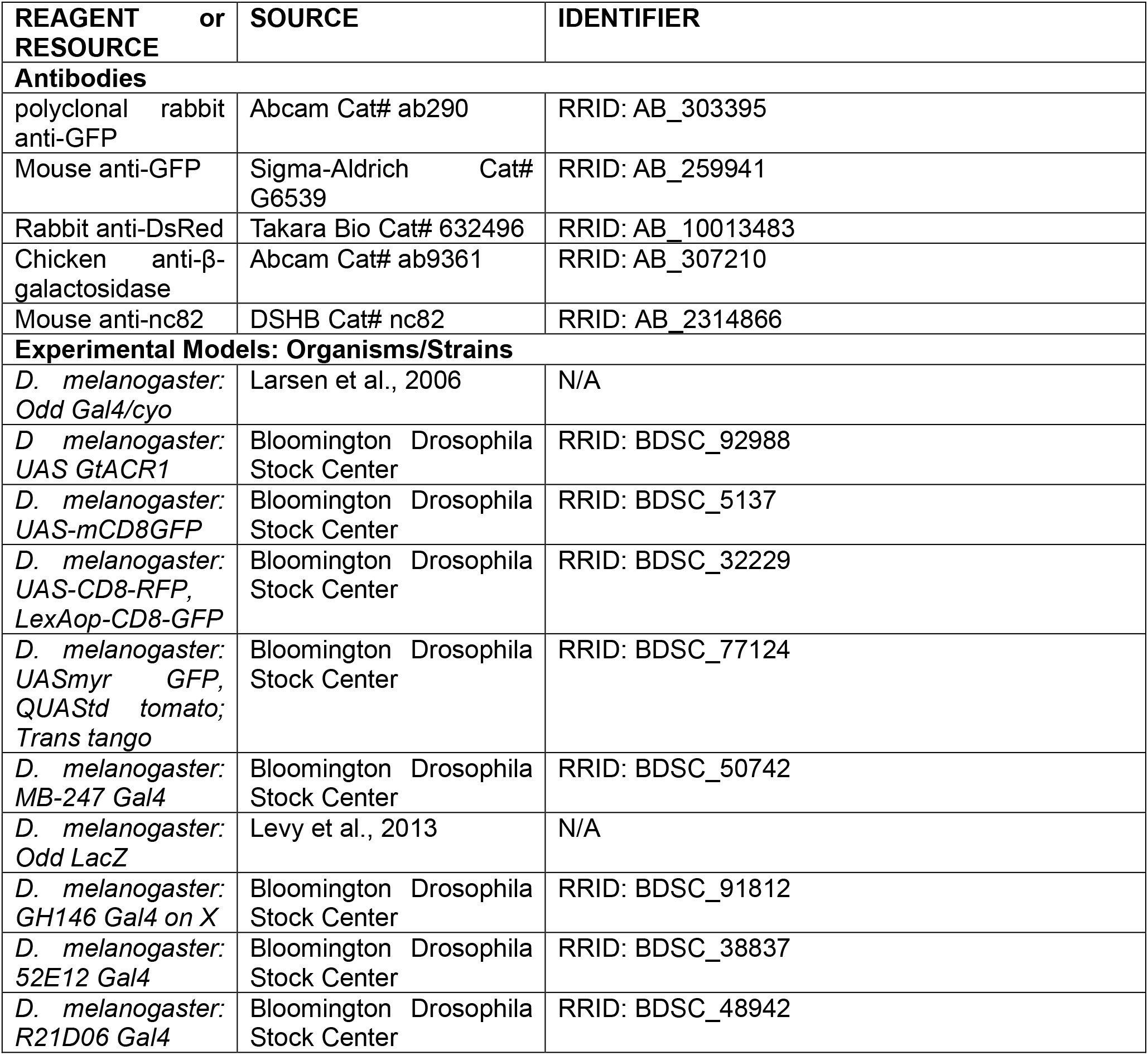
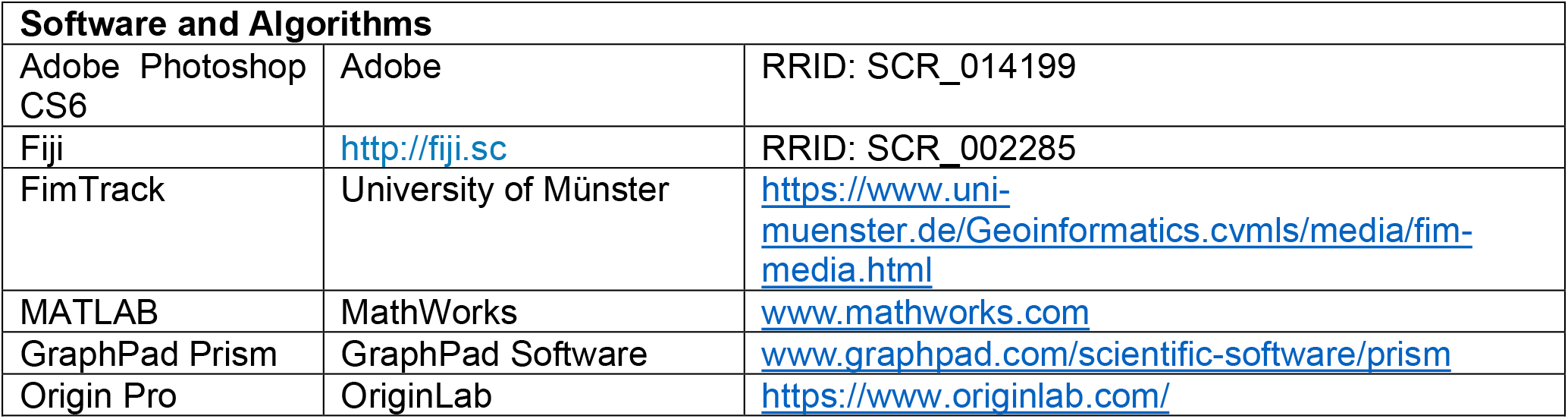

### Key Resources Table

#### Experimental Model and Subject Details

##### Generation and rearing of *Drosophila* stocks

Flies were maintained on a conventional cornmeal-agar molasses medium at 22°C and 60– 70% relative humidity, in a 12-hour light/dark cycle.

##### Innate olfactory chemotaxis assay

Chemotaxis assay was performed as described in a previous study^27^. In brief, 80-100 flies were allowed to lay eggs on standard apple juice plates with yeast paste for 12 h at 25^°^C. The plates were then incubated for a further 3 days at 25^°^C. 24 hours before the experiment, 1 mM all-trans retinal (ATR, R2500; Sigma Aldrich) was added to the plates for optogenetic activation. The plates were covered with aluminum foil to prevent activation of anionic channelrhodopsin GtACR1 from light. Controls were raised and treated in similar conditions. Larvae were collected, washed, and transferred to a 1% agarose layer in a Petri dish, followed by 2-hour starvation. All the assays were subsequently carried out at room temperature in red light conditions. The assay was done on a 90 mm Petri plate with 1% agarose. A 1 cm wide middle was marked as the central zone, serving as the starting point for the larvae. Approximately 30-40 larvae were placed in the center of the plate, separated from each other, before the addition of the odor. The Odor was diluted in paraffin oil (AM 1:1500). 20 μL of the odor solution and paraffin as a control solution were added to the odor cup, which was placed parallel to the central zone where the larvae were located. Optogenetic inactivation was achieved using a green 512nm LED light source.

After introducing the odor to the arena, the lid was placed on a Petri plate, and larvae were allowed to wander for 5 minutes. After 5 minutes, the number of larvae was counted on each side to calculate the Response Index (RI) as follows:

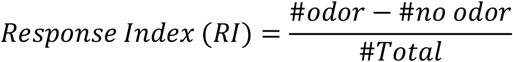

A positive RI means attractive chemotaxis and a negative RI means repulsive chemotaxis. RI with 0 indicates no preference for the odor over the control side.

##### Immunohistochemistry

Larval brain dissections were conducted in 1X phosphate-buffered saline (PBS), followed by fixation in freshly prepared 4% paraformaldehyde (PFA) in 1X PBS for 30 minutes at room temperature. Post-fixation, the specimens underwent four successive washes with 0.3% PTX (0.3% Triton X-100 in 1X PBS) for 15 minutes each at room temperature. Subsequently, a 15-minute blocking step at room temperature was performed using 0.1% PBTX (0.1% normal goat serum in 0.3% PTX).

For immunostaining, primary antibodies were diluted in 0.1% PBTX and incubated with the samples on a shaker at 4°C for 12 hours. Subsequent washes in 0.3% PTX (15 minutes each, repeated four times, totaling 60 minutes) followed this incubation step.

Secondary antibodies, conjugated with Alexa Fluor 488, 568, and 647 (1:500; Invitrogen), were diluted in 0.1% PBTX and applied to the samples for a 2-hour incubation at room temperature on a shaker. After removing the secondary antibody, further washing steps were executed using 0.3% PTX (15 minutes each, repeated four times, totaling 60 minutes).

The imaging procedure was conducted utilizing a Zeiss 510 confocal microscope. Z-stacks were systematically acquired with optical sections set at 1-μm intervals. Subsequently, the raw image data were exported to NIH ImageJ for comprehensive image analysis. Additional image processing was performed using Adobe Photoshop.

##### Associative learning assay

Odor-fructose associative learning was performed as described in a previous study^24^. Briefly, larvae were reared under conditions similar to those employed for the innate chemotaxis assay. Two cohorts of 25-30 larvae each underwent a training regimen involving exposure to an odor and fructose as a reward, either in paired or unpaired groups.

For paired group training, larvae were placed in a petri dish containing 1% agarose mixed with a 6% fructose solution, and n-Isoamyl acetate (AM; diluted 1:1500 in paraffin oil) for 5 minutes. Subsequently, the larvae were transferred to another petri dish filled solely with 1% agarose for an additional 5 minutes. This sequence was repeated three times.

Similarly, in the unpaired group training, larvae were placed in a petri dish containing 1% agarose and AM for 5 minutes, then transferred to another petri dish with a 6% sucrose solution. This cycle was also repeated three times.

Post-training, the larvae were positioned in the center of a Petri plate filled with 1% agarose, with AM odor on one side and paraffin oil on the other. The larvae were allowed to roam freely for 3 minutes. Optogenetic inactivation was performed using a green light source as needed. Larvae were then counted on each side to compute the response index. After computing the response index for paired and unpaired groups, we calculated the performance index as follows:

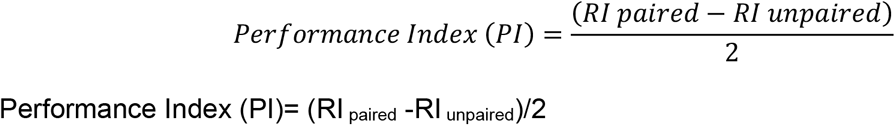

These performance indices range from -1 to 1. A positive value indicates appetitive learning, while a negative value indicates aversive learning.

For baseline behavior after training, larvae were tested for response index in the presence of reward.

To examine appetitive associative memory with two odors, larvae were divided into two groups. One group was trained with AM (1:1500 dilution) paired with a reward, while the other was trained with undiluted OCT paired with the same reward. Post-training, their Response Index (RI) was tested on agarose plates with AM and OCT at opposite ends.

The association between odor and aversive stimulus was established using electric shock instead of fructose in the aversive learning paradigm. In the paired group, the AM odor was paired with a 100 V AC for 1 minute, interspersed with a 30-second rest period, repeated for a total duration of 5 minutes. Conversely, for the unpaired group, the presentation of odor and electric shock followed a separation protocol similar to that of the unpaired training experiment involving fructose.

##### Live imaging and locomotion tracks

Larval behavior was captured using a Nikon Digital Sight DS-Ri1 camera at 5 frames per second (fps) during chemotaxis and associative learning assays. The recorded videos were analyzed using FIMTrack for behavioral assessment. Subsequently, data from FIMTrack were analyzed using customized MATLAB programs designed to calculate turn rates, following a methodology adapted from Gomez Marin et al^23^.

##### Statistical analysis and graph

GraphPad Prism 9 was used for all statistical analyses. Each assay was repeated 10 times, and statistical significance was determined using non-parametric statistics (Kruskal–Wallis test and Mann–Whitney U-test). Graphs and images were created using Origin Pro.

